# Ancestral function of Inhibitors-of-kappaB regulates *Caenorhabditis elegans* development

**DOI:** 10.1101/2020.03.26.010355

**Authors:** David Brena, Joan Bertran, Montserrat Porta-de-la-Riva, Yolanda Guillén, Eric Cornes, Dmytro Kukhtar, Lluís Campos-Vicens, Lierni Fernández, Irene Pecharroman, Albert Garcia-López, Khademul Islam, Laura Marruecos, Anna Bigas, Julián Cerón, Lluís Espinosa

## Abstract

Mammalian IκB proteins (IκBs) exert their main function as negative regulators of NF-κB, a central signaling pathway controlling immunity and inflammation. An alternative chromatin role for IκBs has been shown to affect stemness and cell differentiation. However, the involvement of NF-κB in this function has not been excluded. NFKI-1 and IKB-1 are IκB homologs in *Caenorhabditis elegans*, which lacks NF-κB nuclear effectors. We found that *nfki-1* and *ikb-1* mutants display developmental defects that phenocopy mutations in Polycomb and UTX-1 histone demethylase, suggesting a role for *C. elegans* IκBs in chromatin regulation. Further supporting this possibility *(i)* we detected NFKI-1 in the nucleus of cells; *(ii)* NFKI-1 and IKB-1 bind to histones and Polycomb proteins, *(iii)* and associate with chromatin *in vivo*, and *(iv)* mutations in *nfki-1* and *ikb-1* alter chromatin marks. Based on these results, we propose that ancestral IκB inhibitors modulate Polycomb activity at specific gene subsets with an impact on development.

## Introduction

The mammalian inhibitors of NF-κB, known as IκBs, consist of several homologues whose main function is the cytoplasmic retention of the NF-κB transcription factors, thus leading to suppression of the pathway under non-activating conditions. Stimuli that activate the NF-κB pathway induce IκB phosphorylation at specific serine (S) residues (32 and 36 for IκBα) imposed by the IKK complex of kinases. This process leads to ubiquitination of lysines (K) 21-22 by the E3 ubiquitin ligase β-TRCP, proteasomal degradation of the repressor, and release of the NF-κB factors. Free NF-κB is then translocated to the nucleus to drive transcriptional activation of specific target genes (Zhang, et al., 2017).

Alternative nuclear roles for IκBα and IκBβ homologues have previously been described thus expanding the influence of this family of repressors beyond NF-κB regulation. SUMO2/3 modification of IκBα impairs its association with the NF-κB factors (Culver, et al., 2010). We identified SUMOylated IκBα as the IκBα variant (with molecular weight of 60 kDa in contrast with the 37 kDa of canonical IκBα) capable of binding chromatin to regulate transcriptional activity of various genes that are important during embryonic development such as *Hox* and *Irx*, in cooperation with elements of the Polycomb Repressor Complex (PRC2) (Mulero, et al., 2013a; Mulero, et al., 2013b). Importantly, stimulation with TNFα led to activation of IκBα-bound genes in an NF-κB-independent manner, associated to the chromatin dissociation of the PRC2 subunits EZH2 and SuZ12. Similarly, Drosophila *Cactus*, the orthologous of mammalian IκB proteins, functionally interacts with components of PRC as indicated by the synergistic homeotic transformation imposed by Cactus and Polycomb mutations that was demonstrated as *Dorsal* (NF-κB) independent (Mulero, et al., 2013b). Nevertheless, exploring the functional role of chromatin-bound IκBα has been hampered by the prominent role of IκB proteins in the regulation of NF-κB pathway, and in the inflammatory and immune responses.

*Caenorhabditis elegans* lacks recognizable NF-κB factors such as RelA, RelB or c-Rel, which in all NF-κB-proficient organisms represent the functional effectors of the pathway. Nevertheless, *C. elegans* contains two homologues for the IκB proteins called NFKI-1 and IKB-1 of estimated molecular weights of 66.5 kDa and 66.4 kDa, the IKK family kinase IKKE-1 (Sato, et al., 2018) and the β-TRCP homologue LIN-23. NFKI-1 is predominantly present in the cytoplasm of neuronal cells and acts downstream of interleukin 17 (IL-17) in a signaling pathway to regulate oxygen sensing (Chen, et al., 2017). IKB-1 has a predominant expression in the cytosol of muscle cells (Meissner, et al., 2011), and *ikb-1* mutants do not exhibit any obvious phenotype (Pujol, et al., 2001) but are less resistant to *S. enterica*-mediated killing (Tenor and Aballay, 2008). Thus, *nfki-1* and *ikb-1* are not essential genes and there are no indications in the literature suggesting nuclear functions for the corresponding IκB proteins. Here we report low penetrance developmental phenotypes in *nfki-1* and *ikb-1* mutants that phenocopy deficiencies in chromatin-related genes encoding the demethylase UTX-1(Agger, et al., 2007; Hong, et al., 2007) and Polycomb proteins (Bender, et al., 2004). We demonstrate that *C. elegans* IκBs are contained at low levels in the nucleus of the cells, they interact with histones and PRC2 elements *in vitro*, and associate with chromatin *in vivo*. *nfki-1* and *ikb-1* mutants have a deregulated transcriptome associated with altered distribution of the chromatin marks H3K27me3 and H3K36me3. Thus, our study indicates the existence of ancestral nuclear functions for IκBs in an animal that lacks NF-κB factors.

## Results

### *nfki-1* and *ikb-1* are the homologs of mammalian IκBs and display specific expression patterns during *C. elegans* development

Phylogenetic trees from the TreeFam project (Schreiber, et al., 2014) located *nfki-1* on the IκBδ, and IκBϵ branch, and *ikb-1* closer to IκBα and IκBβ (**Figure S1A** **and** **B**). However, direct analysis of protein sequence similarity using BLASTP identified NFKI-1 and IKB-1 as the most likely orthologs of human IκBα and BCL3, respectively (**Figure 1A** **and** **1B**). BCL3 was initially identified in mammals as an IκB family member involved in the regulation of p50-NF-κB-mediated gene transcription (Franzoso, et al., 1992; Kerr, et al., 1992). The region with the highest sequence similarity between NFKI-1 and IKB-1 includes a series of ankyrin repeats, which are known to mediate the interaction between IκBs and the NF-κB transcription factors (Jacobs and Harrison, 1998). Interestingly, NFKI-1 but not IKB-1 contains the IKK phosphorylation consensus sequence (D<u>S</u>GXX<u>S</u>), which is exclusive of the IκB family of repressors and highly conserved in the mammalian IκBα, IκBβ and IκBϵ homologues, and the *Drosophila* protein Cactus (**Figure 1C**). Western blot analysis using a monoclonal antibody against p-S32 of human IκBα from EGFP∷NFKI-1 immunoprecipitates (endogenously expressed in *C. elegans* L4 stage) confirmed the presence of a phosphorylated NFKI-1 at the IKK consensus site (**Figure 1D)**. Lysine (K) 21, localized 12 aminoacids upstream of the phosphorylation domain of IκBα, which is either ubiquitinated to initiate proteasomal degradation, or SUMOylated to impose nuclear IκBα function (Mulero, et al., 2013b; Culver, et al., 2010; Desterro, et al., 1998), is also conserved in NFKI-1 (**Figure 1C**). We identified additional SUMOylation consensus sites at the N-terminal half of both NFKI-1 and IKB-1, and several putative SUMO-interaction regions using the GPS-SUMO tool (Zhao, et al., 2014; Ren, et al., 2009) (**Figures S1C** **and** **S1D**).

**Figure 1.**
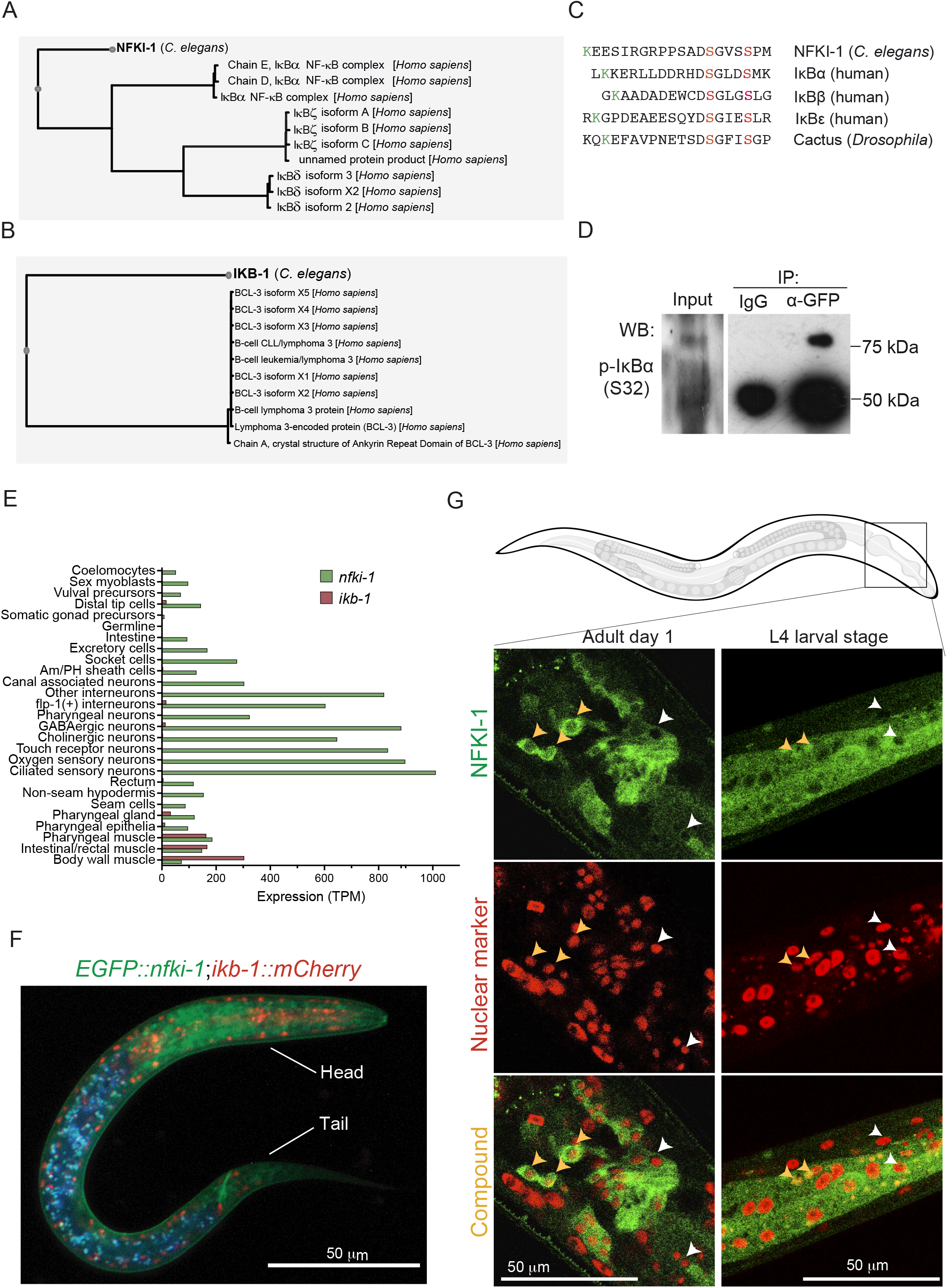
Homology of NFKI-1 and IKB-1 with human IκB proteins and cellular and subcellular patterns of expression. **(A, B)** Image of the first-hit alignment of the *C. elegans* NFKI-1 (A) and IKB-1 (B) proteins obtained with the BlastP program (Camacho, et al., 2009). Data for (A) and (B) were retrieved from GExplore 1.4 (www.genome.sfu.ca/gexplore) (Hutter and Suh, 2016). **(C)** Alignment of different IκB proteins, including NFKI-1, centered on the region that contain the consensus phosphorylation sites (residues 32 and 36 for IκBα, in red) for IKKβ kinase. Note the conservation of the upstream lysine residue that is either ubiquitinated or SUMOylated (in green) in IκBα. **(D)** Western blot analysis using the anti-p-S32-IκBα of *C. elegans* protein lysates of endogenous *EGFP∷nfki-1* strain (L4 stage) precipitated with α-GFP antibody. IgG precipitation is included as specificity control. **(E)** Tissue-specific expression profiles of L2 animals (Cao, et al., 2017). *nfki-1* (green) is mainly expressed in the nervous system and *ikb-1* (red) in the muscular system. Expression data is presented as transcripts per million reads (TPM). **(F)** Microscopy image of an L1 animal carrying both EGFP∷NFKI-1 and IKB-1∷mCHERRY endogenous reporters. DAPI channel is merged to indicate gut autofluorescence. Scale bar: 50 μm. **(G)** Confocal microscopy images of transgenic animals showing the expression pattern of EGFP∷NFKI-1 at the indicated developmental stages. mCHERRY∷SFTB-1 is used as nuclear marker. Yellow arrowheads denote cells that show a nuclear and cytoplasmic EGFP∷NFKI-1 signal. White arrowheads point cells that do not display a nuclear but only cytoplasmic localization at the same plane. Scale bars: 50 μm. Imaging was done with 63X magnification in Z-stacks with 0.25 μm of distance between planes. Images represent a single plane.

We searched in public transcriptomic datasets for the expression patterns of *nfki-1* and *ikb-1* in different tissues and developmental stages. Temporal series of RNA-sequencing (RNA-seq) during *C. elegans* embryonic development showed that *nfki-1* and *ikb-1* are expressed in late embryos, coinciding with morphogenesis and final differentiation of embryonic cells, and at all larval stages with elevated expression at dauer stage, which is acquired in starvation conditions (Boeck, et al., 2016; Hutter and Suh, 2016; Gerstein, et al., 2010) (**Figure S2A).** A dataset of Single cell Combinatorial Indexing RNAseq (sci-RNAseq) at L2 stage shows that *nfki-1* expression is rather ubiquitous in somatic cells but enriched in neuronal cells, whereas *ikb-1* is mainly restricted to muscle cells (**Figure 1E**) (Cao et al, 2017; Hutter and Suh, 2016).

To compare the cell type distribution of IKB-1 and NFKI-1, we characterized CRISPR-engineered endogenous fluorescent reporters and observed a low overlap between *nfki-1* and *ikb-1* postembryonic expression patterns as predicted by transcriptomic studies. At L1 stage, *EGFP*∷*nfki-1* expression was clearly detected in neuronal cells of the head and tail (**Figure 1F**). To explore the possibility of *nfki-1* being expressed at low levels in other cell types, we produced a multicopy transgene that overexpressed *nfki-1* under the control of its own promoter (several strains with extrachromosomal or integrated as low copy after gene bombardment). These transgenic animals displayed a ubiquitous *nfki-1∷TY1∷EGFP∷3xFLAG* expression and an evident subcellular location in both cytoplasm and nuclei (**Figure S2B**). We also investigated the subcellular location of NFKI-1 by generating a strain that contains endogenous EGFP∷NFKI-1 and a reporter for a nuclear protein (the splicing factor SFTB-1) (Serrat et al, Plos Genetics 2019). By confocal microscopy, we observed that NFKI-1 was predominant in the cytoplasm, but a low percentage of cells displayed nuclear NFKI-1 distribution (**Figure 1G**). Similarly, IKB-1:mCHERRY was detected predominantly in the cytoplasm in a subset of cells (**Figure S2C).** These results fit with the predominant cytoplasmic distribution of IκBs in most mammalian cells (see Human Protein Atlas at www.proteinatlas.org; (Uhlen, et al., 2010)) and the reported function of NFKI-1 associated to cytoplasmic signaling pathways (Chen, et al., 2017), but are also compatible with *C. elegans* IκB orthologs playing specific nuclear functions.

### Stress response and developmental defects in *nfki-1* and *ikb-1* mutants

Mutants for *nfki-1* and *ikb-1* were previously characterized without any obvious developmental defect despite their altered responses to cytokine signaling (Chen, et al., 2017; Pujol, et al., 2001). By CRISPR-Cas9, we created additional mutant strains to avoid phenotypes due to side mutations on the existing strains (produced by mutagenesis) (**Figure 2A**). First, since *nfki-1* and *ikb-1* are both upregulated under dietary restriction (Ludewig, et al., 2014), we conducted a survival assay with starved L1 *nfki-1* and *ikb-1* mutants to explore their sensitivity to environmental stresses. We found that *nfki-1(cer2)*, *ikb-1(cer9)* and the double mutants survived longer in the absence of food compared with both the sensitive mutants and wild type worms (**Figure 2B**).

**Figure 2.**
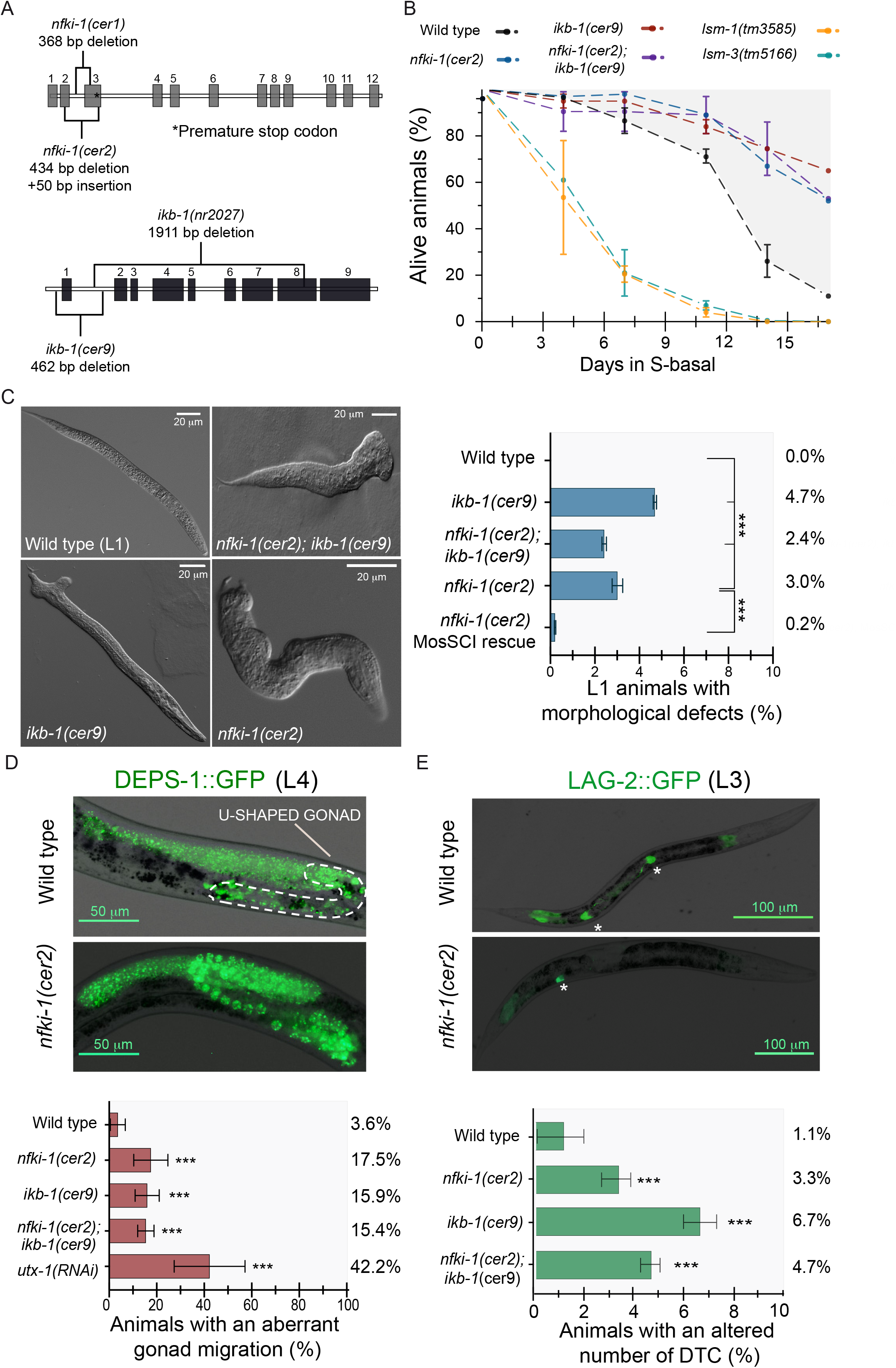
Phenotypes of *nfki-1* and *ikb-1* mutants. **(A)** Graphical representation of the *nfki-1* and *ikb-1* mutants used in the study. *cer1* allele is a CRISPR-generated 368 bp deletion near *nfki-1* exon 2. *cer2* is a 434 bp deletion and 50 bp insertion at exon 2. Both modifications generate premature stop codons. *cer9* allele is a 462 bp deletion that includes the ATG start codon of *ikb-1. nr2027* is a 1911 bp deletion of *ikb-1* that includes the ankyrin-repeat domains encoding sequence (Pujol, et al., 2001). **(B)** Survival curves of the indicated nematode genotypes growing under nutrient restriction. Mutants for the stress proteins LSM-1 and LSM-3 are shown as sensitive controls (Cornes, et al., 2015). n=100 animals per time point. N=2. Error bars show standard deviation (SD) of 2 experiments. Statistically significant differences were calculated using two-tailed Spearman correlation test (CI: 95%). **(C)** Diferential Interference Contrast (DIC) images showing the different morphological defects observed in *nfki-1* mutant animals. The graph indicates the percent of larvae with aberrant morphology, which is rescued after *nfki-1* wild-type copy insertion by MosSCI in chromosome II (Chen, et al., 2017). n>3000. N=2. Error bars show SD of 2 experiments. Statistically significant differences were calculated using two-sided Chi-square test (CI: 95%). **(D)** Representative microscopy images of wild type and *nfki-1(cer2)* worms displaying gonad migration defects. The dashed white line depicts the typical U-shape of the normal gonad in the image of the wild type animal. Graph shows the percentage of animals with an aberrant gonad migration phenotype. n>60. N=3. Error bars show SD of 3 experiments. Statistically significant differences were calculated using two-sided Chi-square test (CI: 95%) **(E)** Alterations in the number of DTCs were visualized using the LAG-2∷GFP reporter line. Note the presence of two DTCs in the WT and a single DTC in the *nfki-1* mutant (white asterisks). Scale bar: 100 μm. Graph display the percentage of animals with an abnormal DTC number. n>1000. N=3. Error bars show SD of 3 experiments. Statistically significant differences were calculated using two-sided Chi-square test (CI: 95%).

Remarkably, a low but significant percentage of *nfki-1(cer2)* and *ikb-1(cer9)* mutants displayed severe morphological defects (**Figure 2C**). To confirm that these defects were caused by IκB mutation*s*, we performed two additional experiments: *(i)* we produced an additional *nfki-1* allele, removing the whole *nfki-1* coding sequence (**Figure S3A**), and observed a similar phenotype (**Figure S3B)** and *(ii)* we rescued the *nfki-1(cer2)* phenotypes with a single copy of *nfki-1* inserted in another chromosome (Chen, et al., 2017) (**Figure 2C**). Of note, these morphological defects are similar to those observed in mutants of *Hox* genes (Zhao, et al., 2010; Van Auken, et al., 2002; Van Auken, et al., 2000) and Polycomb mutants (Capowski, et al., 1991), suggesting that IκB deficiency affected PRC2 function, as we described in *Drosophila* and the mammalian skin and intestine (Marruecos, et al., 2020; Mulero, et al., 2013b). We also observed consistent defects in gonad migration of *nfki-1(cer2)* and *ikb-1(cer9)* mutant animals, which are analogous to those found upon RNAi depletion of the H3K27 demethylase gene *utx-1* (**Figure 2D**), also producing severe morphogenetic defects at L1 (Vandamme, et al., 2012). An aberrant gonad migration could be consequence of a defective Distal Tip Cell (DTC), which is a somatic cell that controls this migration process (Wong and Schwarzbauer, 2012). Since single cell transcriptomics of *C. elegans* larvae detected expression of *nfki-1* and *ikb-1* in DTCs (**Figure 1E**), we studied the cellular differentiation of these cells by using the LAG-2∷GFP marker. A small percentage of animals (3% to 6%) displayed an aberrant number of DTCs (mostly one DTC instead of two) (**Figure 2E**). This result suggests that the aberrant gonad phenotype observed in *nfki-1* and *ikb-1* mutants could be due to defects in specification or differentiation of the DTCs. Interestingly, a double mutant strain for *nfki-1* and *ikb-1* did not increase the phenotype, suggesting that these genes may act in the same genetic pathway.

Altogether, our results indicate that IκB proteins contribute to the regulation of *C. elegans* development, probably through modulation of Polycomb activity, that impacts on the differentiation state of specific cell types.

### NFKI-1 and IKB-1 physically interact with histones and PRC2 proteins *in vitro* and bind to the chromatin *in vivo*

Supporting a role for *C. elegans* IκBs in chromatin regulation, by pull down assays, we demonstrated that HA-tagged NFKI-1 and IKB-1 expressed in mammalian HEK-293T cells bind histone H2A and H3, and to a minor extent H4 (**Figure 3A**). We then determined whether NFKI-1 and IKB-1 bind the *C. elegans* PRC2 subunits MES-2, MES-3 and MES-6. Both NFKI-1 and IKB-1 bound PRC2 elements *in vitro*, and the interaction domain was specific for each PRC2 protein (**Figures 3B-E**). In contrast, neither NFKI-1 nor IKB-1 expressed in mammalian cells interacted with p50 or p65 NF-κB proteins in co-precipitation experiments (**Figure 3F**).

**Figure 3.**
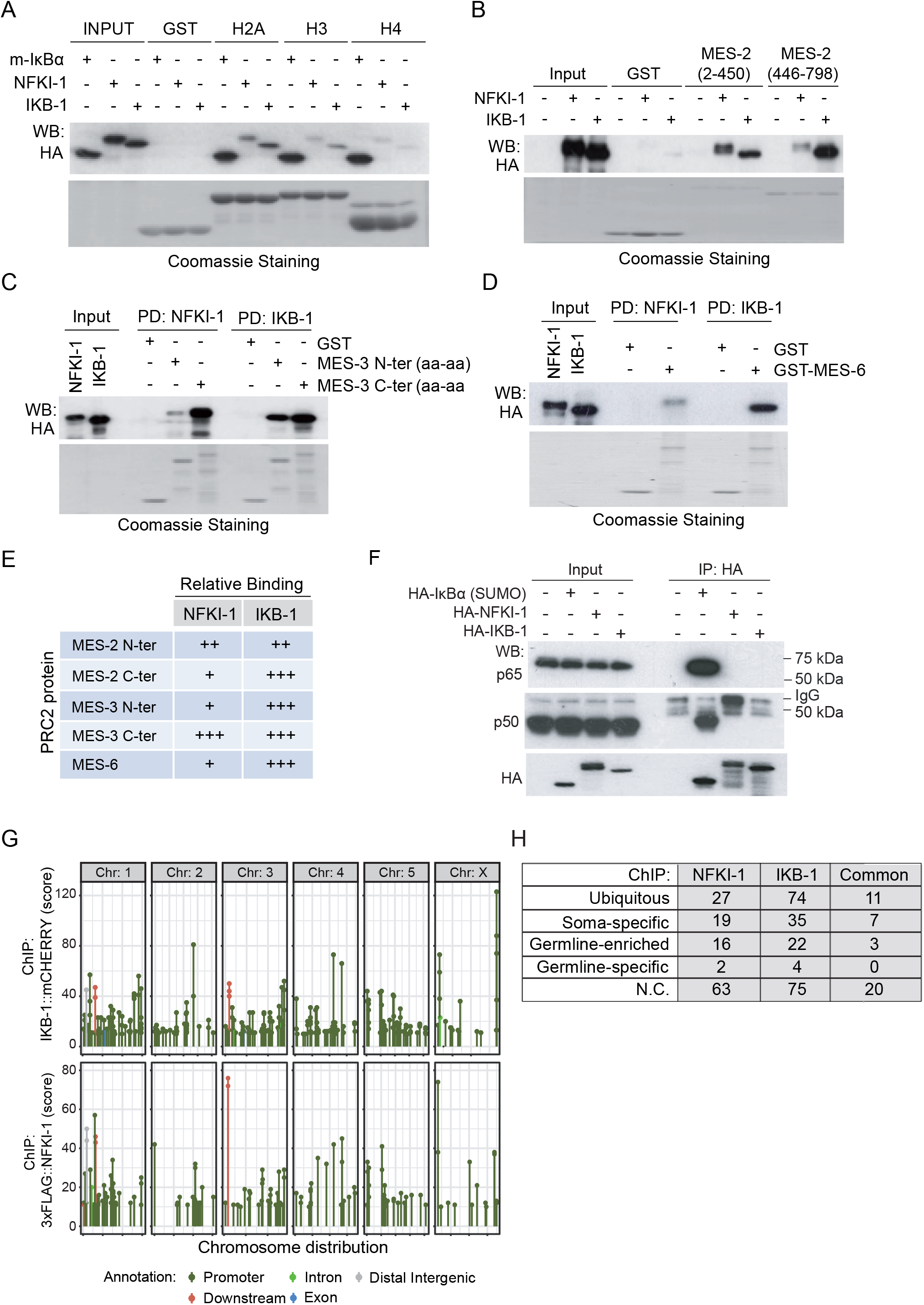
*C. elegans* IκBs physically interact with histones and PRC2 proteins, but not mammalian NF-κB proteins *in vitro* and bind chromatin *in vivo*. **(A)** Pull-down assay using HA-tagged NFKI-1, IKB-1 or mammalian SUMO-IκBα expressed in HEK-293T cells and the indicated GST-fusion histone constructs as bait **(B-D)** Pull-down assays from HA-tagged NFKI-1 or IKB-1 containing extracts using GST-fusion proteins containing the indicated fragments of MES-2 **(B)**, MES-3 **(C)** or MES-6 **(D)**. **(E)** Summary of the data shown in B-D. **(F)** Extracts from HEK-293T cells transfected with HA-tagged human SUMO-IκBα or *C. elegans* NFKI-1 and IKB-1 were used in co-immunoprecipitation experiments to measure association with the NF-κB proteins p65/RelA and p50. Western blot analysis of a representative experiment is shown. **(G)** Distribution of peaks from 3xFLAG∷NFKI-1 and IKB-1∷mCHERRY ChIP-seq across *C. elegans* chromosomes, indicating the localization of the peaks relative to the closest annotated gene. Data are presented relative to input. **(H)** Table indicating the cell-type category distribution of genes identified in the ChIP analysis.

Next, we performed chromatin immunoprecipitation followed by sequencing (ChIP-seq) in 3xFLAG∷NFKI-1; IKB-1∷mCherry endogenous reporters at L1 stage using anti-FLAG and anti-mCherry antibodies. We detected association of NFKI-1 or IKB-1 to chromatin at 127 and 210 genes, respectively. NFKI-1 and IKB-1 ChIP peaks primarily localized at the promoter regions of genes and were similarly distributed among chromosomes (**Figure 3G**) (**Table S1)**. Among potential IκB target genes we identified soma-specific, germline-enriched and ubiquitously expressed, and none of these categories was notably predominant (**Figure 3H**).

Together these results indicate that NFKI-1 and IKB-1 have the capacity to bind chromatin, likely through direct interaction with histones, as well as other chromatin-related proteins such as PRC2 subunits.

### Gene expression changes in IκB-deficient mutants correlate with altered Polycomb-deposited chromatin marks

To validate the possibility that NFKI-1 and IKB-1 exert specific functions in the chromatin of *C. elegans* cells, we synchronized *C. elegans* larvae at L4 stage of wild type and mutant strains, and worm extracts were split in two groups to perform RNA-seq and ChIP-seq experiments with equivalent samples. We identified a large cohort of differentially expressed genes (DEG) in *nfki-1(cer1)* and *ikb-1(nr2027)*. Interestingly, there was a huge correlation between DEG in both mutants (**Figure 4A**) (**Table S2**), suggesting that NFKI-1 and IKB-1 regulate a similar set of genes, most likely in distinct cell types. Among the subset of DEGs, we identified a germline-specific signature that was enriched among upregulated genes in both mutants (**Figure 4B**). We performed RNA-seq of *nfki-1* and *ikb-1* mutants at L1 stage and we did not observe upregulation of germline genes (**Figure S4A**), but we still detected a correlation between DEGs in both mutants (**Figure S4B**). Principal component analysis (PCA) of RNA-seq data clustered together L1 WT and IκB mutants, in contrast with L4 WT samples that were separated from *nfki-1* and *ikb-1* mutants, indicating that transcriptional regulation imposed by IκB during postembryonic development results in the most distinct transcriptome at L4 (**Figure S4C**).

**Figure 4.**
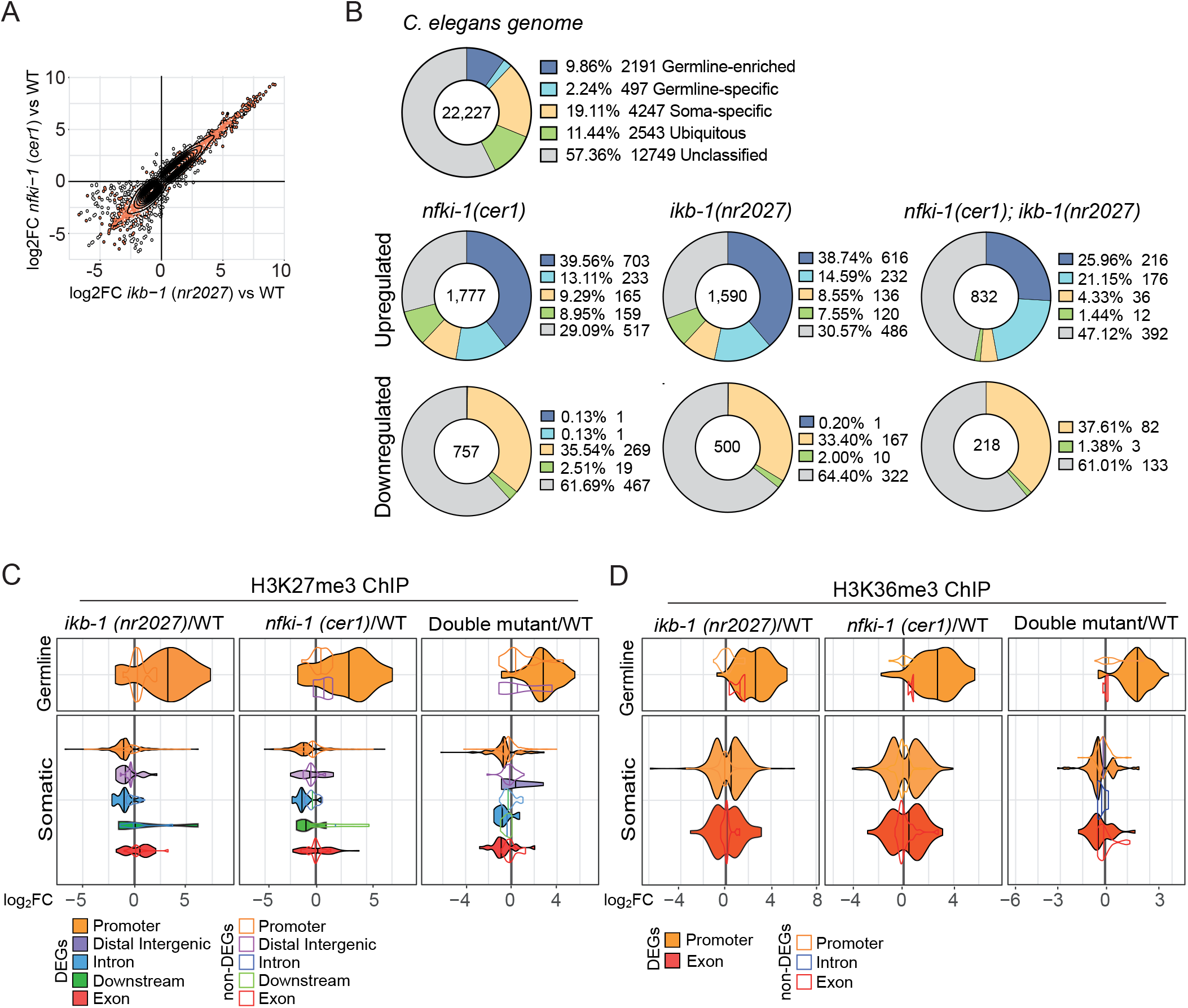
RNA-seq and ChIP-seq indicate that gene expression and distribution of regulatory chromatin marks are affected in IκB-deficient mutants. **(A)** 2D plot illustrating the correlation between genes differentially expressed (adjusted p-value < 0.01, log_2_ fold-change ≥ 2) in *nfki-1* and *ikb-1* deficient animals at L4 larvae stage. R=0.947, p-value < 2.2e-16, Spearman test. **(B)** Doughnut charts showing the distribution of genes categorized according to whether their expression is ubiquitous (green), germline-enriched (dark blue), germline-specific (light blue), soma-specific (yellow) or unclassified (gray) for differentially expressed genes at L4 stage. Numbers in the center represent the number of genes in each dataset. Categories dataset was extracted from (Gaydos, et al., 2012). Statistically significant differences between expected and observed distribution was calculated using Chi-square test for goodness of fit (p<0.05). **(C, D)** Violin plots indicating the density and distribution of H3K27me3 (C) and H3K36me3 (D) peaks at germline and soma-specific genes differentially (color-filled/black lines) and not differentially (colored lines) expressed in the indicated genotypes (p-adj < 0.05). x-axis represents the expression levels (log_2_ fold-change) in mutants relative to the wild type.

Then, we explored whether differential expression of genes observed in IκB mutants was the result of a general alteration in the PRC2-imposed chromatin mark H3K27me3 (Ahringer and Gasser, 2018). Western blot analysis indicated that *nfki-1* and *ikb-1* mutants showed comparable levels of the repressive H3K27me3 mark (**Figure S4D**). Then, we performed ChIP-seq analyses for H3K27me3 and the activating mark H3K36me3, which is mainly antagonistic to H3K27me3 (Ahringer and Gasser, 2018), on the L4 paired samples used for RNA-seq. Most of the ChIP-seq peaks were located at promoter regions and displayed a similar distribution among the different IκB deficient backgrounds (**Table S1**). A more detailed analysis of the data indicated that in soma-specific DEGs, the H3K27me3 mark was accumulated at the promoter and intronic regions of genes downregulated in *nfki-1(cer1)*, *ikb-1(nr2027)* and the double mutant, but randomly distributed in the non-differentially expressed group (non-DEG) of genes **(Figure 4C)**. In germline genes, which are generally up-regulated in the mutants, both H3K27me3 and H3K36me3 marks were found only in promoter regions and presented a high accumulation of H3K36me3 mark (**Figure 4C** **and** **4D**).

Overall, our results here demonstrate an ancestral function of NFKI-1 and IKB-1 as regulators of key chromatin marks deposition and transcriptional activation of a subset of genes with impact on *C. elegans* development. This is the first evidence of transcriptional regulation imposed by IκB homologues in the absence of a functional NF-κB pathway.

## Discussion

IκB proteins are mainly known because of their function as suppressors of inflammation and repressors of NF-κB. We have taken advantage of a non-NF-κB system to discriminate between NF-κB-dependent and-independent functions of these essential proteins. The presence of highly conserved IκB proteins in an NF-κB-deficient organism is at least surprising, based on the fact that free IκBα is intrinsically unstable in mammalian cells (half-live of 5-10 minutes) and requires being stabilized through its association with NF-κB (three orders of magnitude prolonged half-live) (Mathes, et al., 2008; O’Dea, et al., 2007; Pando and Verma, 2000).

Importantly, we found that *C. elegans* IκBs are, in fact, unable to bind mammalian p65-NF-κB but they bind PRC2 and histones. The association with these proteins may contribute to their stabilization, mostly dependent on the association to NF-κB in mammalian cells (Mathes, et al., 2008). These results, together with the observation that IκB deficiencies in *C. elegans* result in an altered H3K27me3 distribution, a deregulated transcriptome and morphological defects suggest that nematode IκBs act as modulators of developmental processes, including the animal body patterning, which is also regulated by PRC2, *Hox* genes and the UTX-1 demethylase (Vandamme, et al., 2012; Agger, et al., 2007; Lan, et al., 2007). Another *nfki-1* and *ikb-1* mutant phenotype related to development was the aberrant migration of distal tip cells (DTCs), which also phenocopied the *utx-1* mutants. Altered migration of DTCs is associated with a defective cellular identity of the somatic gonad precursors. Therefore, the body morphology defects as well as the phenotype in DTCs are related to cell fate identity and maintenance, and would be related to nuclear functions of *C. elegans* IκB proteins.

In addition, we found that *nfki-1* and *ikb-1* mutants are more resistant to starvation than wild type animals. This phenotype could be related with the recently demonstrated interaction of NFKI-1 with cytoplasmic MALT-1, which regulate resistance to environmental stresses (Flynn, et al., 2019).

We generated endogenous reporters for *nfki-1* and *ikb-1*. In the case of *nfki-1*, we further studied the subcellular location of NFKI-1, which is predominantly cytoplasmic as previously reported (Chen, et al., 2017). We challenged the reporter strain in different ways, but we did not find a condition, besides overexpression, that increases the presence of NFKI-1 in the nuclei. Supporting the nuclear location of NFKI-1, a recent study detected NFKI-1 in the nuclear fraction of *C. elegans* and a protein-protein interaction with the histone methyl transferase CBP-1/P300 (Flynn, et al., 2019).

The fact that NFKI-1 and IKB-1 deficiencies result in almost identical phenotypes, both at the morphologic, chromatin and transcriptomic levels suggest that both homologues have similar functions in gene regulation. They overlap in a small subset of cells (**Figure 1E**) but, they may exert comparable nuclear functions in distinct cell types to regulate a similar set of genes. Another possibility is that NFKI-1 and IKB-1 deficiency affects either germline development or the speed of larval development thus leading to progressive changes in gene expression, which we detected at L4 stage.

NFKI-1 and IKB-1 present an amino acid stretch highly rich in lysines (K) in the region from aa 31 to 144 (20% of all amino acids). The average K content for non-ribosomal proteins is 5% and 10% for ribosomal proteins (Lott, et al., 2013). So it is plausible to speculate that post-translational modifications of the amino-terminal half of the protein, including ubiquitination and/or SUMOylation, are required for compensating the high excess of positive charge imposed by the high proportion of K residues. Supporting this possibility, we have identified multiple SUMOylation consensuses in both proteins as well as SUMO-binding regions that could be mediating the potential interaction in some cells.

It is still under debate whether NF-κB was originally present in a common ancestor of arthropods and nematodes and then lost in the latter, or NF-κB signaling was originated in a common ancestor of arthropods and vertebrates after the divergence of nematodes (Kim and Ausubel, 2005). Thus, our results suggest two possibilities: *(i)* a nuclear function of ancestral IκBs has emerged in an NF-κB-free scenario and has primarily been maintained by BCL3 in mammals, whereas IκBα has functionally diverged to inhibit NF-κB, maintaining the capacity to regulate a fraction of PRC2-repressed genes (when phosphorylated and SUMOylated) or *(ii)* NF-κB pathway was originally present in *C. elegans* being primarily regulated by NFKI-1, and subsequently lost, thus rewiring NFKI-1 toward its moonlighting function as regulator of PRC2 target genes. In any case, from the evolution point of view, according to the cytoplasmic roles of *C. elegans* NFKI-1 and IKB-1, IκB proteins were involved in innate-immunity and stress responses even before the existence of NF-κB.

## Material and Methods

### *Caenorhabditis elegans* strains

*C. elegans* strains were cultured on Nematode Growth Medium (NGM) agar plates and maintained by standard methods (Porta-de-la-Riva, et al., 2012; Stiernagle, 2006). We used N2 (Bristol) as wild type strain. All experiments were conducted at 25°C, unless otherwise stated. Before performing assays, strains’ genotype was verified and then animals were grown for at least two generations at the experimental temperature. All strains in this study are listed in **Table S3**. All primer sequences are described in **Table S4**.

### CRISPR-Cas9 editing and nematode transgenesis

For CRISPR-Cas9 editing, single guide RNAs (sgRNAs) were designed using *CCTOP* (Stemmer, et al., 2015) online tool. We followed a co-CRISPR approach (Kim, et al., 2014) using *dpy-10(cn64)* allele as a marker to enrich for successful genome-editing events. Endogenous reporters were obtained using a Nested CRISPR strategy (Vicencio, et al., 2019). PHX267[*ikb-1(syb267[ikb-1∷mCherry*])I] endogenous reporter was provided by Suny Biotech.

Young adult hermaphrodites were injected using XenoWorks Microinjection System. F_1_ progeny was screened by PCR using specific primers and F_2_ homozygotes were confirmed by Sanger sequencing. The sequences of the CRISPR reagents and primers used are available in the **Table S4**.

For the generation of NFKI-1 overexpression reporter lines, fosmid vectors containing an TY1∷EGFP∷3xFLAG-tagged version of *nfki-1* (C33A11.1) were requested from the TransgeneOme resource (Sarov, et al., 2012) and transformation was performed by bombardment with gold particles (Biolistic Helium Gun, Caenotec). *unc-119(ed3)* young adults were shot with 16 μg of the purified DNA of interest (Praitis, et al., 2001). Fosmid constructs were verified by digestion with restriction enzymes. Expression patterns were characterized for 2 independent lines with an integrated transgene and 6 with an extrachromosomal array. All generated lines displayed similar expression patterns.

### RNA isolation

Synchronized animals were collected in M9 buffer after growing for 36 hours (L4s) or 5 hours-fed and 3 hours starved (L1s) at 25°C. Suspended worms were washed 5 times in M9 buffer to remove all traces of bacteria. Total RNA was isolated using TRI Reagent (MRC, Inc.) and PureLink® RNA Mini Kit (Ambion) according to manufacturer’s instructions.

### *C. elegans* microscopy and image processing

Animals were mounted on 2% agar pads with 6 mM tetramisole hydrochloride. Live fluorescence imaging was performed with a Zeiss Axio Observer Z1 inverted fluorescence microscope and confocal imaging was conducted with a Leica TCS SP5 Confocal Laser Scanning Microscope. Imaging was done with 10-63X magnification in Z-stacks with 0.25-1μm distance between planes. We used Zeiss Zen 2012 (Blue Edition), FIJI (ImageJ version 2.0.0-rc-68/1.52p) for image processing, and Adobe Illustrator CS5 for assembly of the figures.

### Phenotypic characterization

For all experiments, gravid adult animals were synchronized by hypochlorite treatment (Porta-de-la-Riva, et al., 2012) and resuspended embryos were allowed to hatch by overnight incubation at 20°C. Then, incubated at 25°C until animals reached the optimal stage for phenotype evaluation.

Larvae morphology was assessed at L1 in a large synchronized population by Nomarski (DIC) microscopy. Animals that presented misshapen bodies, mainly at the posterior region and in the tail, which were completely absent in N2 (Bristol) wild type control, were counted as morphologically aberrant. Gonad migration was characterized in a synchronized population at L4 stage (36h at 25°C) for wild type and mutant backgrounds by DIC and fluorescence microscopy. To facilitate inspection and counting, knockout mutants were crossed with endogenous DEPS-1∷GFP germline reporter. Gonads were considered with an aberrant migration when the expected U-shape was not achieved due to an irregular turn.

Distal tip cell number was examined by microscopy of a synchronous population of wild type and in an IκB-mutant background at L3 stage (24h at 25°C). An abnormal distal tip cell number was considered when animals displayed one or three LAG-2∷GFP-expressing distal tip cells.

### Starvation assay

L1 starvation assay was performed as previously described (Cornes, et al., 2015; Zhang, et al., 2011). Briefly, gravid adult worms were synchronized by hypochlorite treatment and the resulting eggs were resuspended in 4 ml of S-basal without cholesterol. Animals were incubated rotating at 20°C. Larval viability was determined by placing ~100 worms every 3 days onto NGM plates and incubated at 20°C for 2 days. Worms that reached at least L4 stage were marked as survivors and survival rates were calculated. Three biological replicates were used for each timepoint.

### RNA interference (RNAi)

RNAi was performed by feeding placing synchronized L1 worms obtained by hypochlorite treatment on NGM plates seeded with the specific RNAi clone-expressing bacteria at 25°C (Timmons, et al., 2001). We used L4440 empty vector (Fire vector library) as a mock RNAi control. Phenotypic characterization was assessed at different developmental stages of the F_1_ progeny.

### Plasmids

GST fusion proteins were generated using pGEX5x.3 (GE Healthcare) as vector. cDNAs were cloned in frame 3’ from the GST coding-sequences using BamHI and XhoI restriction sites. For full-length proteins, cDNAs were amplified by PCR starting at sequences coding for amino acid 2 and ending at the stop codons. When preparing fusion proteins with particular fragments, appropriate primers were designed to clone in frame and a stop codon was introduced with the reverse primer. Primers were designed with 5’ extensions that included the relevant restriction enzyme recognition sites and additional nucleotides to ensure sensitivity to the restriction enzyme in the PCR product. Sequences used for *mes-2*, *mes-3* and *mes-6* were taken from genebank with accession numbers: NM_064591; NM_001026321 and NM_001026149. For MES-2 and MES-3, two different fusion proteins were constructed to facilitate their production in *E. coli* due to their large size. Thus, for MES-2 the N-terminal construct contains amino acids 2-450 and the C-terminal construct amino acids 446-798. Likewise, for MES-3 the N-terminal construct contains amino acids 2-370 and the C-terminal construct amino acids 321-754. Primer sequences are given in **Table S4**.

Expression plasmids for NFKI-1 and IKB-1 were constructed using the pcDNA4TO vector (ThermoFisher Scientific) previously modified to include two HA-epitope units in the 5’ end. The exact sequence is available upon request. Briefly, cDNAs coding for proteins NFKI-1 (NM_078139) and IKB-1 (NM_060174) were amplified using primers with appropriate 5’ extensions to facilitate cloning by restriction enzyme digestions followed by ligation. Primers are given in **Table S4**.

### Preparation of GST and GST-fusion proteins

Fresh colonies of *E. coli* containing the desired plasmids were used to start fresh cultures for recombinant protein isolation. When the culture OD_600_ reached 0.6, IPTG was added to a final concentration of 0.1M and the culture was further incubated for an additional 3.5 hours at 37°C. Protein expression was verified in SDS-PAGE by Coomassie blue staining. For purification cells were pelleted and resuspended in lysis buffer (10 ml per each 50 ml of bacterial culture; 20mM Tris HCl pH 7.4; 1M NaCl; 0.2 mM EDTA; 1mM PMSF; 1mM DTT; 1mg/mL lysozyme; 1 mini complete tablet (protease inhibitor cocktail Roche), sonicated using 3 cycles of 10 seconds at 25% amplitude and then centrifuged at maximum speed for 30 minutes at 4°C. Supernatant was incubated with glutathione sepharose beads, equilibrated in lysis buffer, for 3 hours at 4°C in a rotary shaker. GST-fusion proteins bound to beads were recovered by gentle centrifugation (1200 rpm for 2 minutes in a microfuge) washes extensively with lysis buffer and finally resuspended in lysis buffer and kept at 4°C. To assess quality and relative quantity of the purified GST-fusion proteins, different volume aliquots were run in SDS-PAGE and stained with Coomassie blue. For the pull-down assay, equal amount of the different GST-fusion proteins was used according to Coomassie blue stain. Additional glutathione-sepharose beads were included when necessary to have a visible bead volume during the process. Beads were equilibrated in eukaryotic lysis buffer (0.5 M Tris HCl pH 7.5 1.5 M NaCl; 10% Nonidet (NP-40); 50 mM EGTA 50 mM EDTA; 200 mM NaF; 1 mini complete tablet/50 ml of buffer (protease inhibitor cocktail; Roche)) and blocked with HEK293T cell extract before use. HEK293T cells overexpressing the proteins of interest were lysed in eukaryotic lysis buffer and insoluble proteins and cell debris were discarded by centrifugation at 13000 rpm for 10 minutes at 4°C. Next, blocked beads were mixed with cell lysate and incubated for 45 minutes at 4°C in a rotary shaker, centrifuged for 2 minutes at 1200 rpm and washed extensively with eukaryotic lysis buffer. Pulled proteins were analyzed by western blot.

### Pull down and peptide co-precipitation assays

*C. elegans* HA-IKB-1 and HA-NFKI-1 expressed in HEK293T cells were used in pull down assays to assess their interaction with histones and members of the Polycomb repressor complex 2. In some experiments, SUMO-IκBα was included as positive control and corresponds to an artificially SUMOylated IκBα variant with a modified (Q90P) SUMO moiety fused to its N-terminus right before amino acid 2. In brief, cells extracts were incubated for 1 hour with the indicated GST or GST fusion proteins bound to glutathione-coated beads. After extensive washing, the presence of HA-tagged proteins bound to the beads was analyzed by western blot analysis.

### Immunoprecipitation

EGFP∷NFKI-1 L4 *C. elegans* were frozen with liquid nitrogen and smashed with mortar and pestle. Then, the cells were lysed with RIPA buffer (0.1% DOC, 10mM Tris-HCl pH 8.0, 140mM NaCl, 1% Triton X-100, 0.1% SDS, 1mM EDTA pH 8.0, 0.5mM EGTA, 10mM NaButyrate, 20mM ◻-Glycerol-phosphate and 100μM NaOrtovanadate), sonicated using 3 cycles of 10 seconds at 25% amplitude and then centrifuged at maximum speed for 30 minutes at 4°C. The proteins in supernatant were precipitated using an antibody anti-GFP [Clontech Ref. 632593]. Rabbit IgG was used as negative control (4μg). The antibody was incubated with lysates overnight and the complexes precipitated with protein A-sepharose beads [GE Healthcare, Ref. 17-0780-01] 2h at 4°C. Precipitated proteins were analyzed by Western blot.

### Western blot

Samples were analyzed by Western blotting using standard SDS–polyacrylamide gel electrophoresis (SDS-PAGE) techniques. In brief, protein samples were boiled in Laemmli buffer, run in polyacrylamide gels, and transferred onto polyvinylidene-difluoride (PVDF) membranes [Millipore Ref. IPVH00010]. Gels were stained with Coomassie Brilliant Blue G-250 [ThermoFischer Ref.20279]. Membranes were incubated overnight at 4°C with the appropriate primary antibodies, extensively washed and then incubated with specific secondary horseradish peroxidase–linked antibodies from Dako [Ref. P0260 and P0448]. Peroxidase activity was visualized using the enhanced chemiluminescence reagent [Biological Industries Ref. 20-500-120] and autoradiography films [GE Healthcare Ref. 28906835].

| Antibody | Company | Reference | Specie | Dilution |
| --- | --- | --- | --- | --- |
| H3K27me3 | Millipore | #07-449 | Rabbit | 1:1000 |
| HA.11 Epitope Tag | Covance | MMS-101R |  | 1:1000 |
| Histone H3 | Abcam | ab1791 | Rabbit | 1:5000 |
| NF-κB p50 | Santa Cruz | sc7178 | Rabbit | 1:1000 |
| NF-κB p65 | Abcam | ab16502 | Rabbit | 1:1000 |
| P-IκBα (S32) | Abcam | Ab92700 | Rabbit | 1:1000 |

### Chromatin immunoprecipitation (ChIP)

Larvae were collected as described by (Askjaer, et al., 2014). Briefly, synchronized animals were grown in *E. coli* OP50-seeded NGM agar 150 mm plates and then collected in M9 buffer after growing for 36 h (L4s) or 5 h-fed and 3h-starved (L1s) at 25°C. Suspended worms (approximately, 200 μl of L1 pelleted animals and 2 ml for L4s) were washed 5 times in M9 buffer to remove all traces of bacteria, resuspended in FA buffer with protease inhibitors cocktail (Roche) and frozen as drops in liquid nitrogen (N_2_).

Then, 1 ml of packed worms was grounded under liquid N_2_ and brought to 10 ml with FA buffer (50 mM HEPES/KOH pH7.5, 1 mM EDTA, 1% Triton X-100, 0.1% sodium deoxycholate, 150 mM NaCl). Next, 600 μl of 37% formaldehyde were added and allowed to crosslink for 20 minutes at room temperature. Glycine (600 μl; 2.5M) was used to quench the reaction, for 2 minutes, and nuclei were recovered by centrifugation (4000 rpm, 5 minutes in a microfuge), washed 3 times with FA buffer by resuspending in 5 ml and spinning again. Finally, nuclei were resuspended in 1.6 ml of FA buffer, split into two Eppendorf tubes and sonicated for 10 minutes in a Bioruptor® Pico (Diagenode) under standard conditions (30s on, 30s off). After sonication, protein extract was quantified and approximately 2 mg of each sample were spun at full speed for 10 minutes at 4°C and the supernatants (SN) were pooled again. SN were pre-cleared with FA-equilibrated SPA/G beads (GE Healthcare) by rotating for 30 minutes at room temperature, recovered by spinning at 1200 rpm for 2 minutes and used for immunoprecipitation (IP). An aliquot of 100 μl was kept as input control at this point. The rest of the sample was split into different tubes for IP with the antibodies of choice. After adding the corresponding antibody (3-10 μg), samples were incubated in a rotator overnight at 4°C and the immunocomplexes were recovered with FA buffer pre-equilibrated SPA/G beads (GE Healthcare) by rotating 2h at 4°C. Beads were pelleted (1200rpm; 2 minutes) washed twice with FA buffer (5 minutes each); once with FA + 0.5 M NaCl (10 minutes); once with FA buffer + 1M NaCl (5 minutes); once with lithium buffer (10 minutes); twice with TE buffer (5 minutes each) and the immunoprecipitated material was recovered by incubation in 100 μl elution buffer (Tris-EDTA; 1% SDS plus inhibitors) 1h at RT. DNA was finally purified by reverse crosslinking at 65°C overnight followed by proteinase K (0.5 or 1 mg/ml, respectively) digestions and purifying with MinElute® PCR purification kit (Qiagen).

| Antibody | Company | Reference | Specie |
| --- | --- | --- | --- |
| Flag M2 | Sigma | F3165 | Mouse |
| mCherry | Invitrogen | PA5-34974 | Rabbit |
| H3K27me3 | Millipore | #07-449 | Rabbit |
| H3K36me3 | Abcam | ab95412 | Mouse |

### ChIP-seq data analysis

DNA samples were sequenced using Illumina® HiSeq platform. Raw single-end 50-bp sequences were filtered by quality (Q>30) and length (length > 20 bp) with Trim Galore (Krueger, 2015). Filtered sequences were aligned against the *C. elegans* genome (WBcel235) with Bowtie2 (Langmead and Salzberg, 2012). MACS2 software (Zhang, et al., 2008); available at https://github.com/taoliu/MACS/) was run first for each replicate, and then combining all replicates, using unique alignments (q-value < 0.1). Broad peaks calling was set for H3K27me3 and H3K36me3 (--broad option). Peak annotation was performed with ChIPseeker package (version 1.10.2) (Yu, et al., 2015).

### RNA-seq data analysis

cDNA was sequenced using Illumina® HiSeq platform, obtaining 30 – 35 million 75 bp single-end reads per sample. Adapter sequences were trimmed with Trim Galore (Krueger, 2015). Sequences were filtered by quality (Q>30) and length (>20 bp). Filtered reads were mapped against *C. elegans* genome (WBcel235) using Bowtie2 (Langmead and Salzberg, 2012). High quality alignments were fed to HTSeq (v.0.9.1) (Anders, et al., 2015) to estimate the normalized counts of each expressed gene. Differentially expressed genes between wild type and knockouts were explored using DESeq2 R package (v.1.20.0) (Love, et al., 2014) considering a threshold of adjusted p-value<0.01.

### Statistical analyses

‘N’ denotes the number of independent replicate experiments performed, while ‘n’ indicates total number of animals analyzed for each condition. Statistical analyses were performed in GraphPad Prism 8 and R. Statistical tests are reported in figure legends. For all graphs, all the p-values were noted according to APA annotation style. p-value>0.05 not significant (n.s.); p-value<0.05 (*); p-value<0.01(**); p-value<0.001 (***).

### Data availability

ChIP-seq and RNA-seq data have been deposited at GEO (GSE146655).

## Acknowledgments

We thank all members of Cerón, Espinosa and Bigas laboratories for helpful discussions and technical support, and Ben Lehner (CRG, Barcelona) for critical reading of the manuscript. This work was funded by grants from Instituto de Salud Carlos III FEDER (PI15/00895, PIE15/00008, PI16/00437 and PI19/00013) and Generalitat de Catalunya 2017SGR135. DB is a predoctoral fellow of the CONACYT “Becas al Extranjero” Program of Mexico. YG is a recipient for CIBERONC contract (Instituto de Salud Carlos III), LM for 2015FI-B00806 and 2016FI-B1 00110, DK for 2018FI_B1_00511, and JG for PERIS SLT002/16/00070 fellowship programs from Generalitat de Catalunya.

## Supplementary Material

### Supplementary Figure legends

**Figure S1.**
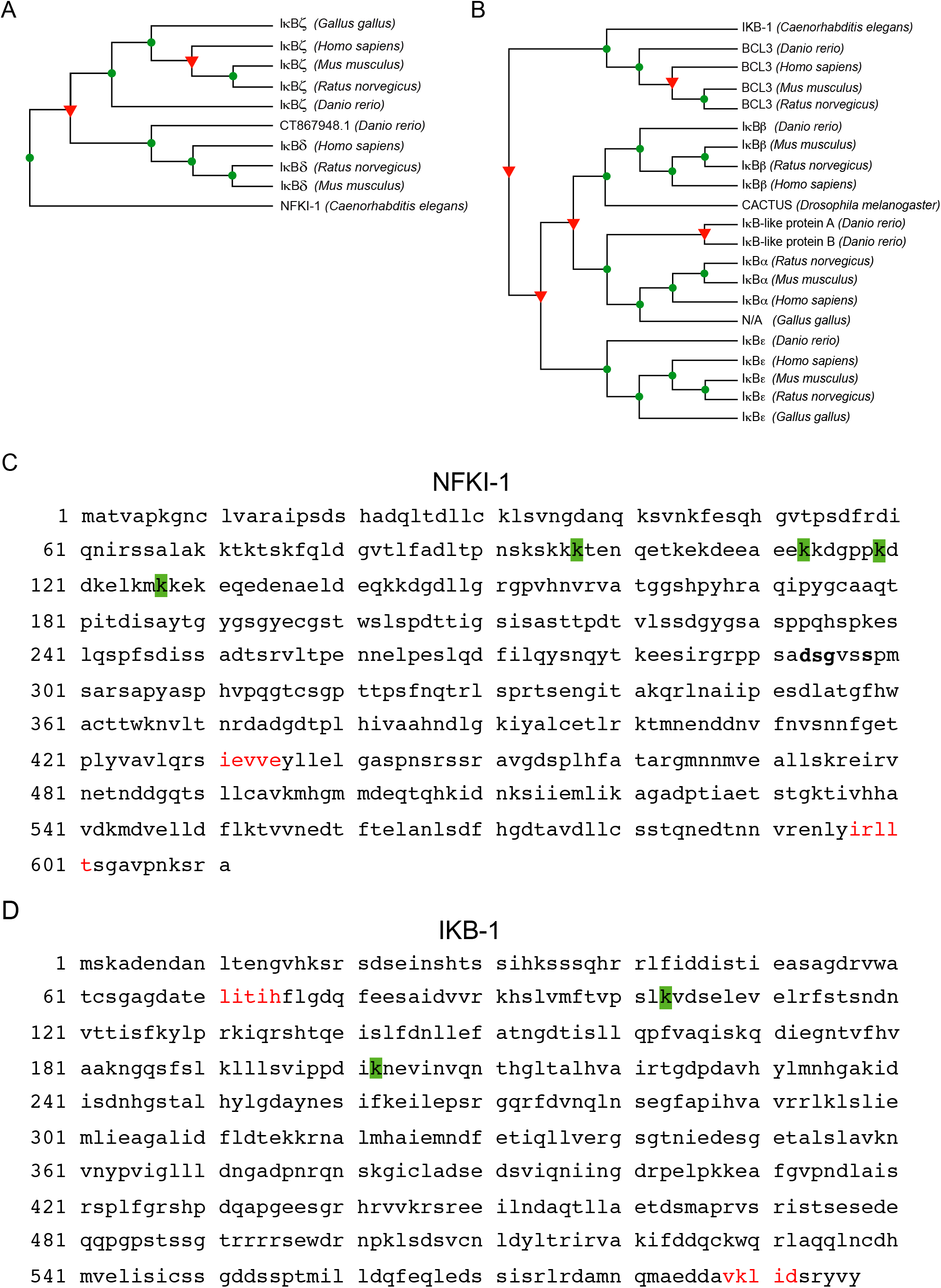
Characterization of NFKI-1 and IKB-1. Related to Figure 1. **(A, B)** Phylogenetic trees for NFKI-1 (A) and IKB-1 (B) proteins. The alignments were 900 and 2305 amino acids long, and presented on average a 52% and 39% of conservation, respectively (Camacho, et al., 2009). **(C, D)** Potential SUMOylated lysines (k, shown in green) and SUMO-interaction sites (in red) in the NFKI-1 (C) and IKB-1 (D) proteins, as predicted using the GPS-SUMO tool (Zhao, et al., 2014).

**Figure S2.**
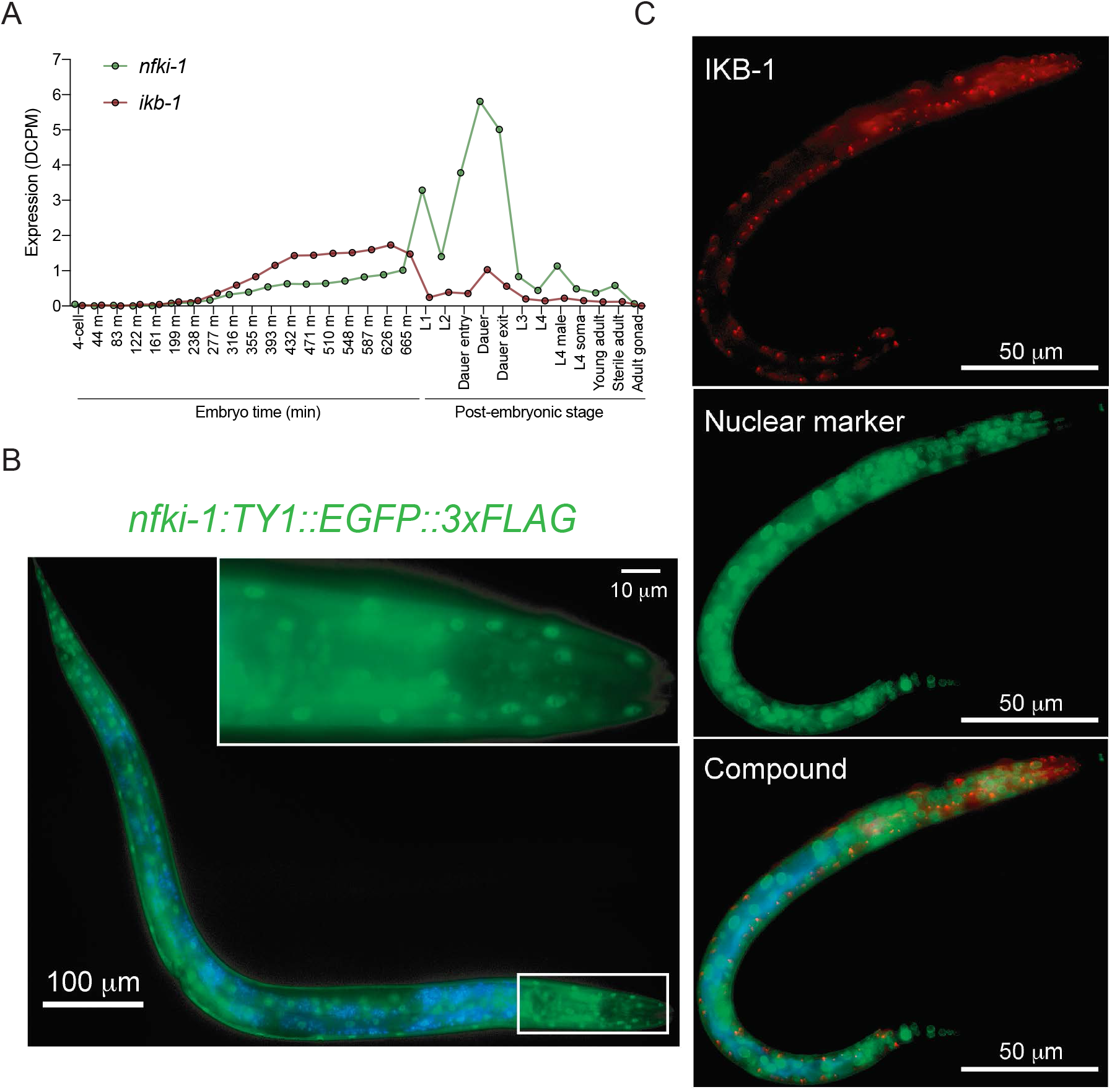
Expression profiles of NFKI-1 and IKB-1. Related to Figure 1. **(A)** RNA-seq time series (Gerstein, et al., 2010) showing *nfki-1* (green) and *ikb-1* (red) stage-specific expression profiles of embryonic development and all postembryonic stages. Expression data is presented as the average depth of coverage per million mapped reads (DCPM). **(B)** Microscopy image of representative CER147 [*cerEx35[nfki-1∷TY1∷EGFP∷3xFLAG+unc-119(+)*] overexpression line at L4 larvae stage. Scale bar: 100 μm. The inset shows the boxed area at higher magnification. Inset scale bar: 10 μm. **(C)** Representative fluorescence microscopy images of a compound IKB-1∷mCHERRY and GFP∷MEL-28 L2 animal. IKB-1∷mCHERRY display dot-like structure signals in the body wall muscle and pharynx tissues. GFP∷MEL-28 endogenous reporter was used as a nuclear envelope marker. DAPI channel was merged to show gut autofluorescence. Scale bars: 50 μm.

**Figure S3.**
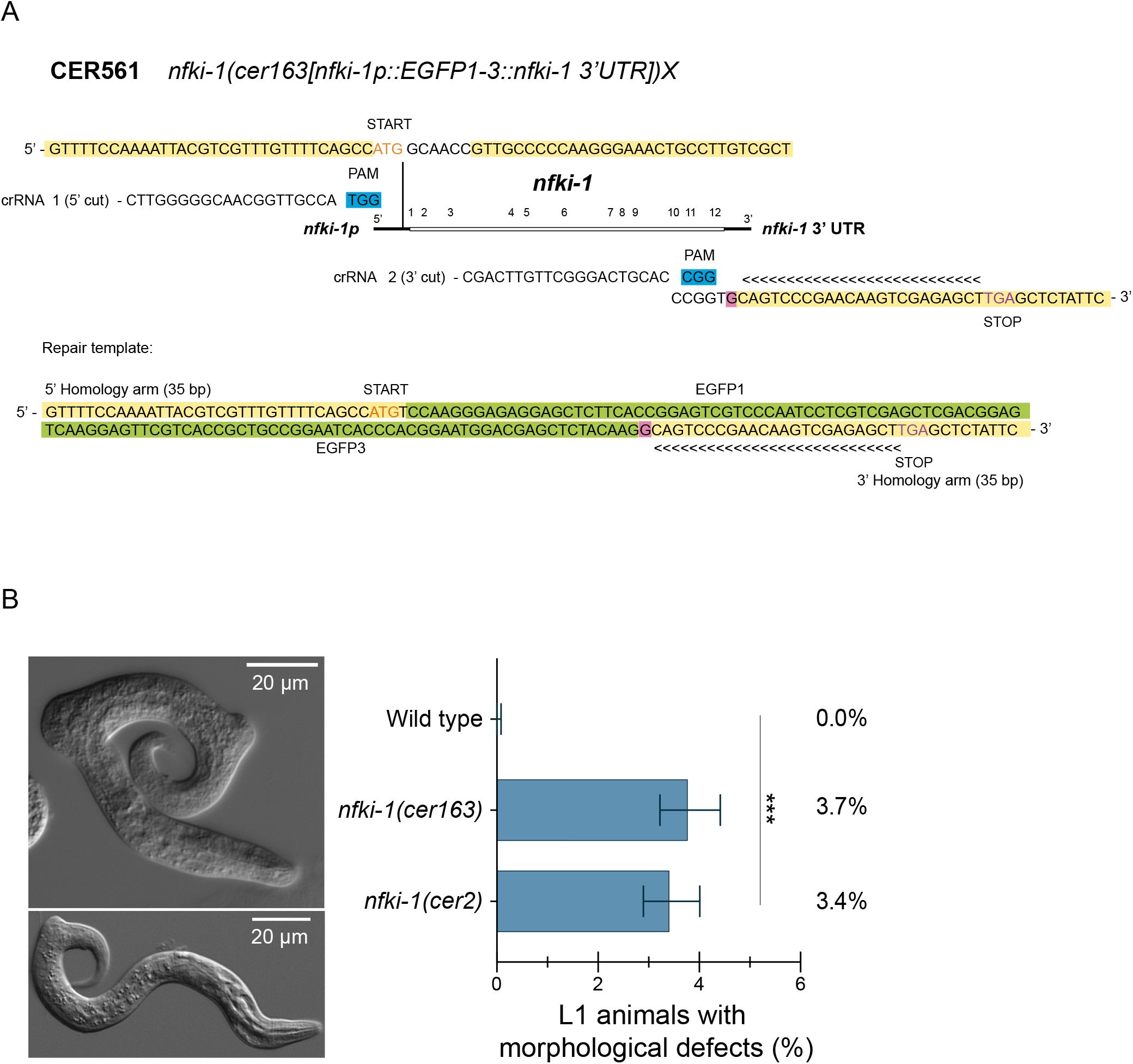
Endogenous reporters’ expression under starvation states and whole CDS deletion allele characterization. Related to Figure 2. (**A**) Schematic representation of whole gene deletion mutant strain CER561[*nfki-1(cer163[pnfki-1∷EGFP1-3])X]* by CRISPR-Cas9. Generation included the use of 2 crRNAs, one near the start codon (1, in orange) and the other near the stop codon (2, in purple). Protospacer adjacent motif (PAM) sequences are depicted in blue and underlined in the sequences. 35 bp-homology arms are highlighted in yellow. Chevron signs indicate *nfki-1*’s CDS last 23 nucleotides, we inserted an additional nucleotide to ensure the preservation of the frame (highlighted in pink). Black arrowheads depict the Cas9 cut site and black arrows show the insertion/deletion site. Full sequence of the repair template is displayed at the bottom. We included the flaking regions of an EGFP in the repair template (termed EGFP1 and EGFP3, respectively, highlighted in green), to facilitate a possible generation of a transcriptional reporter by Nested CRISPR methodology (Vicencio, et al., 2019). **(B)** Representative DIC images showing morphological defects observed in *nfki-1(cer163)* mutant animals. Scale bars: 20 μm. The graph indicates the percent of larvae with aberrant morphology, which is similar to the percentage observed with *nfki-1(cer2)* allele. *n>3500.* N=1. Error bars show upper and lower limits of 95% confidence intervals (CI) calculated by Wilson/Brown method of 1 experiment. Statistically significant differences were calculated using two-sided Chi-square test (CI: 95%).

**Figure S4.**
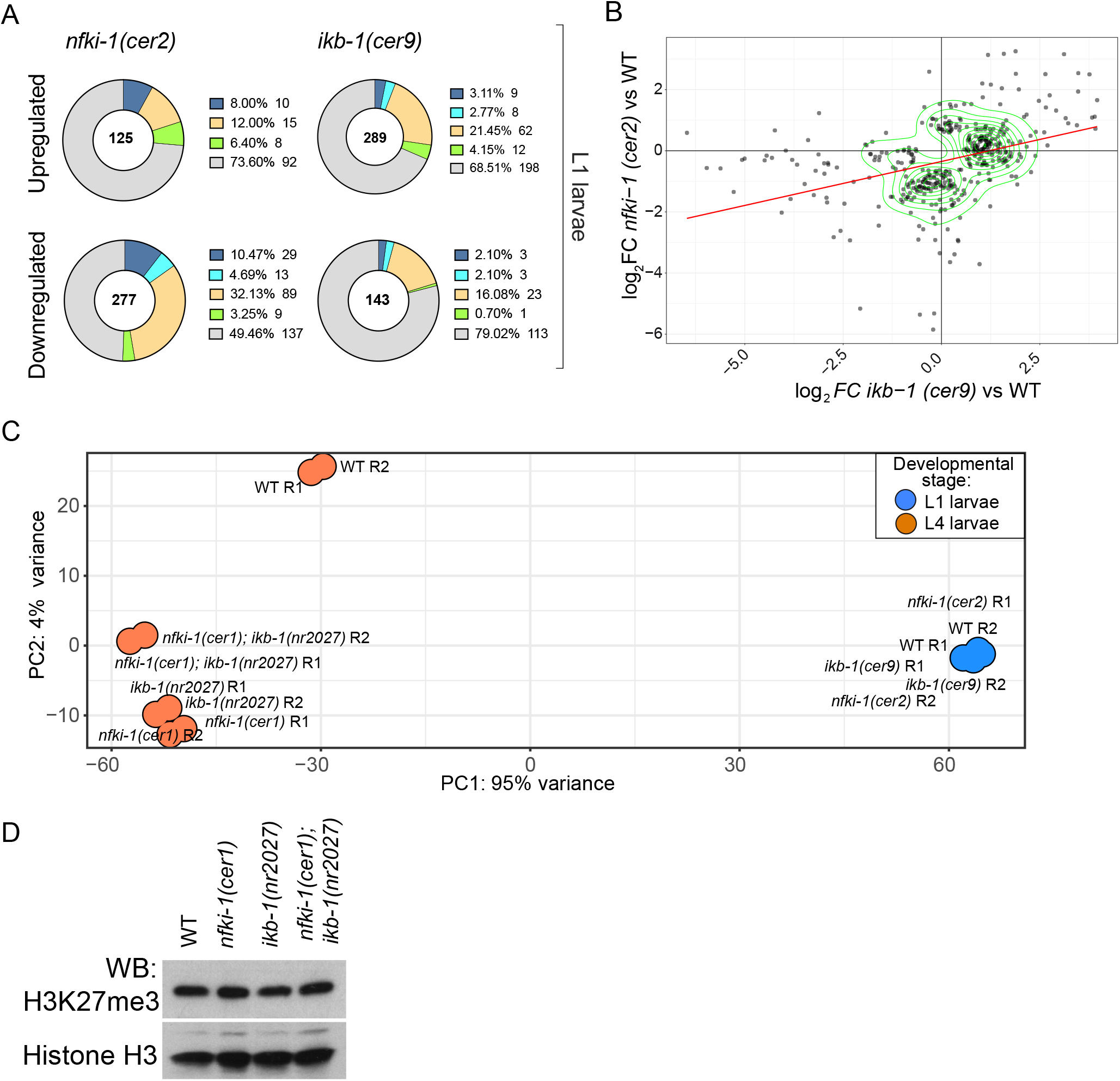
RNA-seqs analyses and H3K27me3 amount in IκB mutants. Related to Figures 3 and 4. **(A)** Doughnut charts showing the distribution of genes categorized according to whether their expression is ubiquitous (green), germline-enriched (dark blue), germline-specific (light blue), soma-specific (yellow) or unclassified (gray) for differentially expressed genes at L1 stage. Numbers in the center represent the number of genes in each dataset. Categories dataset was extracted from (Gaydos, et al., 2012). Statistically significant differences between expected and observed distribution was calculated using Chi-square test for goodness of fit (p<0.05) **(B)** 2D plot illustrating the correlation between genes differentially expressed in *nfki-1* and *ikb-1* deficient animals at L1 stage. **(C)** PCA analysis of RNA-seq samples at L4 (orange) and L1 (blue) for wildtype (WT) and *nfki-1* and *ikb-1* mutants. ‘R’ denotes each replicate. **(D)** Western blot analysis of total protein lysates of L4 worms with the indicated genotypes using an H3K27me3 antibody. Histone H3 is used as a control.

### Supplementary tables

**Tables S1. ChIP-seq analyses.** List of peaks and annotations of 3xFLAG∷NFKI-1 and IKB-1∷mCHERRY ChIP-seq at L1 stage. List of peaks and ChIP-seq annotation of H3K27me3 and H3K36me3 marks in nfki-1 and ikb-1 mutants.

**Table S2. RNA-seq analyses.** List of differentially expressed genes of IκB mutants’ RNA-seq at L4 stage. Classification of DEGs according to (Gaydos, et al., 2012).

**Table S3. List of strains used in this study**.

**Table S4. Primers sequences and CRISPR reagents used in this study**.

